# Expression of tyrosine hydroxylase gene from rat leads to oxidative stress in potato plants

**DOI:** 10.1101/2020.05.15.098210

**Authors:** Kamil Kostyn, Aleksandra Boba, Anna Kostyn, Michał Starzycki, Jan Szopa, Anna Kulma

## Abstract

Catecholamines are biogenic aromatic amines common among both animals and plants. In animals they are synthesized via tyrosine hydroxylation, while in plants, both hydroxylation or decarboxylation of tyrosine are possible, depending on the species, though no tyrosine hydroxylase – a counterpart of animal enzyme has been identified yet. It is known that in potato plants it is the decarboxylation of tyrosine that leads to catecholamine production. In this paper we present the effects of induction of an alternative route of catecholamine production by introducing tyrosine hydroxylase gene from rat. We demonstrate that an animal system can be used by the plant, however, it does not function to synthesize catecholamines. Instead it leads to elevated reactive oxygen species content and constant stress condition to the plant which responds with elevated antioxidant level and further with improved resistance to infection.

**One sentence summary:** Introduction of rat tyrosine hydroxylase gene to potato disturbs catecholamine synthesis, causes oxidative stress and activates antioxidant response.

## Introduction

Catecholamines are natural biogenic amines found both in animals and plants. Their biosynthesis route in animals has been well recognized. Shortly, it starts with tyrosine hydroxylation to L-DOPA, catalyzed by tyrosine hydroxylase (TH, EC: 1.14.16.2). L-DOPA is further decarboxylated to dopamine by L-DOPA decarboxylase (DD, EC: 4.1.1.28). This neurotransmitter may undergo hydroxylation by dopamine β-monooxygenase (DH, EC: 1.14.17.1) to norepinephrine, and further to epinephrine by phenylethanolamine N-methyltransferase (PNMT, EC: 2.1.1.28). This main route of catecholamine biosynthesis may be complemented by that employing tyramine as a substrate for dopamine biosynthesis in a reaction driven by a cytochrome P450 isoform (CYP2D6 in human brain or CYP2D in rats) (Wang et al., 2014). Functions of catecholamines in animals (hormones, neurotransmitters) are well recognized, as they have been studied extensively for decades (Goldstein, 2010).

Interestingly, the presence of these compounds in plants was shown as early as in the fifties of the previous century (Udenfriend et al., 1959) and confirmed in a number of plant species since (Kulma and Szopa, 2007). The role of catecholamines in plants is not fully recognized. Numerous publications report either on their structural or regulatory functions. They may be substrates for other relevant compounds, like betalains (Gandia-Herrero and Garcia-Carmona, 2013), alkaloids, like mescaline (Aniszewski, 2015), melanins or hydroxycinnamic acid amides. Catecholamine conjugates with hydroxycinnamic acids are important molecules produced upon pathogen infection that may be incorporated into the cell wall reinforcing its structure and providing a tighter barrier for the invading microorganisms (Newman et al., 2001). Their role may be also explained by their antioxidant properties, since catecholamine molecules are potent free radical scavenging agents (Kostyn et al., 2012). This activity is even more pronounced in their amide conjugates. Due to neighboring hydroxyl groups in the catechol ring, catecholamines possess remarkable ability to quench free radicals (Kanazawa and Sakakibara, 2000). The DPPH and superoxide scavenging activities of catecholamines were shown to be much higher than those of ascorbic acid and comparable to those of catechin and flavan (Rahman et al., 2006). Moreover, catecholamines reacted with single oxygen distinctively stronger than catechin or sodium azide (Shimizu et al., 2010). Dopamine was shown to be a stronger antioxidant than tocopherol and glutathione (Kostyn et al., 2012), and flavonoids – luteolin and quercetin in vitro (Huang et al., 2019). However, it must be noted that oxidized forms of catecholamines might be toxic and can pose threat to cellular components. Analogously to their function in animals, catecholamines were suggested to control carbohydrate metabolism. In transgenic potatoes they influenced simple sugar content with simultaneous decrease in starch (Swiedrych et al., 2004, Skirycz et al., 2005). Moreover, catecholamine interaction with phytohormones was reported. Catecholamines were shown to stimulate ethylene biosynthesis (Dai et al., 1993) or induce flowering in *Lemna paucicostata* (Yokoyama et al., 2000). Dopamine was indicated as a mediator of gibberellic acid activity in lettuce hypocotyl (Kamisaka, 1979) and inhibited auxin oxidase in tobacco and *Acmella oppositifolia* leading to growth stimulation (Protacio et al., 1992).

Studies on catecholamine biosynthesis pathway in plants showed their analogy to that found in animals. However, while in animals the route through tyrosine hydroxylation is preferred, in plants both hydroxylation and decarboxylation of tyrosine was confirmed, though not simultaneously, and the active route is probably species-specific (Kulma and Szopa, 2007). Tyrosine decarboxylase (TD) presence in plants has been established and its structure and functions well recognized (Zhu et al., 2016). Plant TDs show narrow substrate specificity and can catalyze decarboxylation of tyrosine only or tyrosine and L-DOPA, depending on plant species and/or tissue and developmental stage (Lehmann and Pollmann, 2009, Świędrych et al., 2004). However, despite the fact that the presence of L-DOPA was evidenced in numerous plants (Soares et al., 2014), tyrosine hydroxylase was not found in these organisms. Hydroxylation of tyrosine may be ascribed to other enzymes. For instance, tyrosinase catalyzes the reactions leading to melanin formation, namely the ortho-hydroxylation of monophenols to o-diphenols (monophenolase activity, EC 1.14.18.1) and the subsequent oxidation of o-diphenols to the corresponding o-quinones (diphenolase activity, EC 1.10.3.1). The L-DOPA produced in this pathway is utilized in the synthesis of DOPA-quinone rather than catecholamines (Chung et al., 2015). However, L-DOPA may be decarboxylated to tyramine with the aid of tyrosine decarboxylase, which further is processed to yield catecholamines (Kulma and Szopa, 2007). Nevertheless, tyrosine hydroxylase activity was found in callus cultures of *Portulaca grandiflora* and it was shown to be different from tyrosinase and polyphenol oxidase activities (Yamamoto et al., 2001). In another study an enzyme of tyrosine hydroxylase activity was purified and characterized in *Mucuna pruriens* (Luthra and Singh, 2010). It was highly specific to tyrosine, but showed also a slight activity to tyramine converting it to dopamine, although catalysis of such reaction is usually ascribed to tyrosinases (Kahn et al., 1998).

L-DOPA, being the precursor for catecholamines can also serve as a substrate for melanins, producing leucodopachrome and dopachrome either by auto-oxidation or due to polyphenol oxidase (PPO, EC 1.10.3.1) activity. This reaction produces reactive oxygen species (ROS), such like hydrogen peroxide, superoxide anion and hydroxyl radical (Soares et al., 2014). L-DOPA, like other non-protein amino acids are accumulated in plants to serve as deterrents to herbivores (Fürstenberg-Hägg et al., 2013). Moreover, it can act as allelochemical and after exudation from the roots leads to reduction of growth of neighboring plants (Fujii et al., 1991). The toxicity of L-DOPA is clearly multi-source. The already mentioned production of quinones is connected with ROS generation on one hand, but it also leads to quinoprotein formation on the other. These proteins can be produced by reaction between dopaquinone and the sulfhydryl groups of proteins and lead to such negative effects like enzyme deactivation, mitochondrial dysfunction, DNA fragmentation, and apoptosis (Mushtaq et al., 2013, Jana et al., 2011). In addition, it has been shown that L-DOPA may be incorporated to proteins via mimicking tyrosine or phenylalanine in respective tRNA synthesis (Rodgers and Shiozawa, 2008). Nevertheless, the ROS generated due to L-DOPA oxidation turn out to be the most serious problem for a plant as they cause oxidative stress with a number of diverse repercussions. Therefore, a specialized machinery of ROS neutralization must be employed by the plants to cope with oxidative stress. This machinery is based on two main groups of agents, one employing enzymatic apparatus to neutralize reactive radicals and another using small antioxidant molecules. The enzymatic mechanisms include antioxidant enzymes, such as SOD (which converts O_2_^.-^ to H_2_O_2_), catalases and peroxidases (which remove H_2_O_2_). The non-enzymatic mechanisms of ROS removal include antioxidant molecules, such as ascorbic acid, glutathione, phenylpropanoids, carotenoids, and α-tocopherol. Reactive oxygen species are intrinsic element of numerous stress events and, acting as signaling molecules, trigger signal transduction pathways in plant response (Huang et al., 2019). It is well known that ROS is the factor that precedes activation of SA signaling in plants responding to biotic and abiotic stresses (Herrera-Vásquez et al., 2015). Members of the TGA and WRKY transcription factor families involved in SA-mediated transcriptional regulation play significant role in controlling biosynthesis of antioxidants, for instance phenylpropanoids, in response to stress.

In this study we show the effects of rat tyrosine hydroxylase gene expression in potato. We show that an induction of the alternative route of tyrosine metabolism toward catecholamines results in the seizure of the catecholamine biosynthesis. L-DOPA synthesized due to TH activity undergoes oxidation with simultaneous ROS production, which generates a state of constant oxidative stress that forces the plant to counteract with activation of antioxidant machinery.

## Results

### Construction of transgenic potato plants

For transgenic potato plants generation agrotransformation method was employed and resulted in 232 regenerants, which were further pre-selected for TH cDNA presence in genomic DNA by means of PCR (Supplementary Fig. S1A). In 118 individuals a positive signal was detected. Plants positively screened for the presence of the transgene in genomic DNA were submitted to a subsequent stage of selection – the assessment of the level of the introduced gene’s expression with use of Northern Blotting technique. Total RNA isolated from 3-week-old potato plants from in vitro culture was separated in gel electrophoresis and then transferred onto nitrocellulose membrane. Among 118 samples hybridized with the radioactive probe in 28 of them a strong signal was detected (Supplementary Fig. S1B). The 28 plants selected by means of Northern blotting analysis were analyzed for the presence of tyrosine hydroxylase protein. Total proteins were separated in acrylamide gel electrophoresis (SDS-PAGE) and transferred onto nitrocellulose membrane, followed by incubation with antibodies specific to *R. norvegicus* tyrosine hydroxylase. Out of 28 plants 18 were positively selected (Supplementary Fig. S1C) for the presence of 56 kDa protein specifically recognized by anti-HT antibodies and based on the results from transcript level analysis and Western blotting 5 transgenic lines were selected for further analyses (HT8, HT12, HT28, HT43 and HT53).

### Metabolic profile of the transgenic potato plants

To determine the influence of the rat tyrosine hydroxylase gene expression on metabolites their contents were measured in methanolic extracts of selected transgenic lines (HTZ lines) grown in in vitro culture by means of GC-MS technique. A number of primary and secondary metabolites were identified. The obtained data were submitted to Independent Component Analysis (ICA). The HTZ transgenic lines clearly separated from the control, and considering both, the IC01 and IC02 components line HTZ28 was the most different from the control (Fig. 1).

**Fig. 1.**
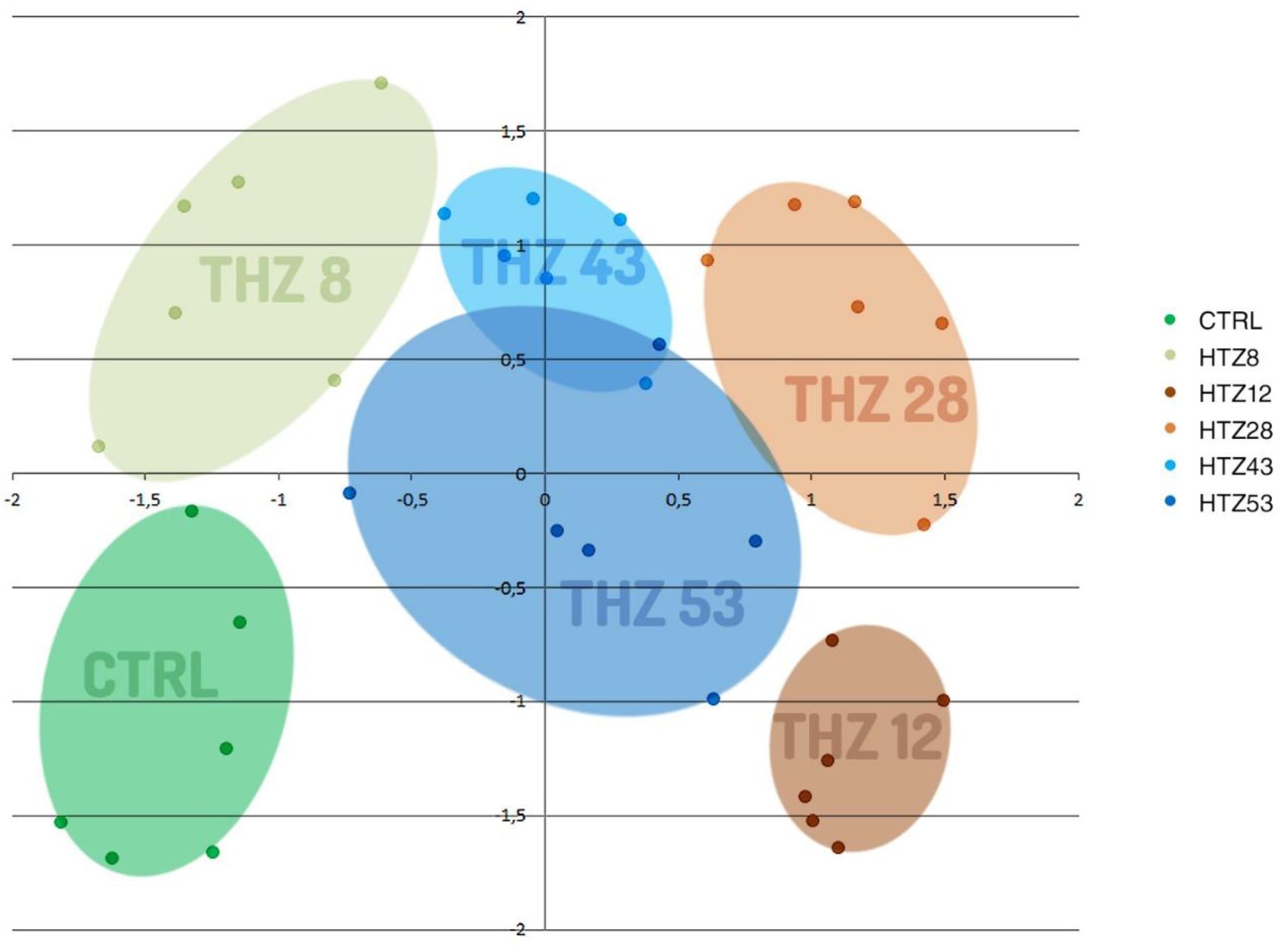
Independent Component Analysis on data obtained from GC-MS analysis of transgenic HTZ lines and the control. Colored areas cover agglomerations of the same sample’s datapoints.

#### Catecholamine level in the transgenic HTZ lines

Levels of the metabolites of catecholamine group were measured in the HTZ lines relatively to the non-transgenic control (Fig. 2). In all of the transgenic HTZ lines the level of tyrosine, a substrate for tyrosine hydroxylase was considerably lower compared to the control. The level of L-DOPA, a direct product of tyrosine hydroxylase was higher in all the transgenic lines reaching 157% of the control. The level of tyramine did not change significantly in the HTZ lines (only in line HTZ43 it was by 30% lower than in the control). Noticeable decrease in dopamine level was measured in the transgenic lines to 10%-48% of the control. Hydroxylation of dopamine at C-2 yields norepinephrine which is further methylated to epinephrine. The level of the latter was lower in all transgenic lines but HTZ8 and the changes reached 62%-82% of the control. The level of metanephrine, a catabolite of epinephrine was lower in all HTZ lines and reached 36%-69% of the control. Homovanillic acid is a product of dopamine degradation. Its level was decreased in the transgenic lines to 36%-74% of the control.

**Fig. 2.**
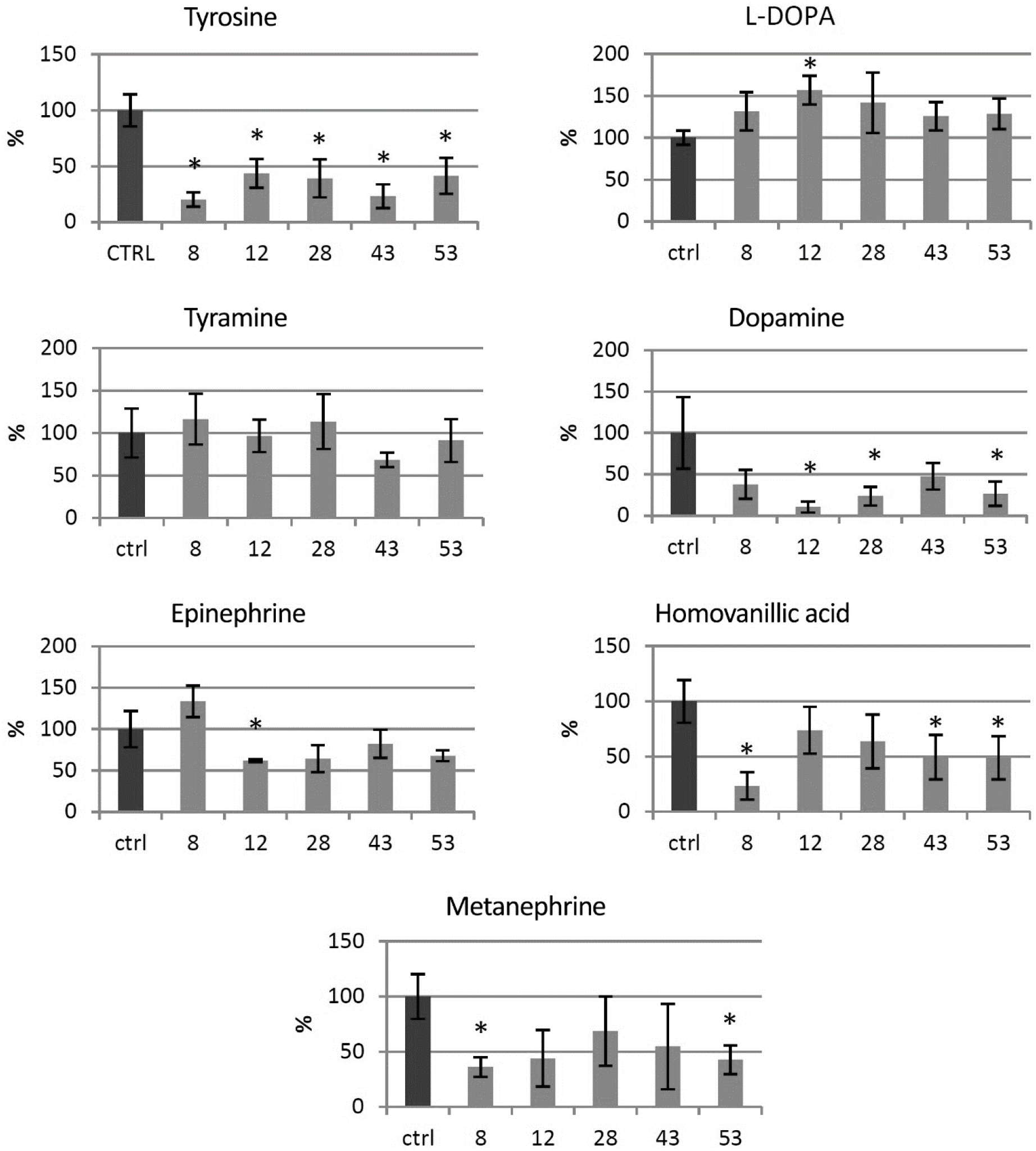
Levels of catecholamine group metabolites in HTZ transgenic lines relatively to the control (non-transgenic potato) obtained with GC-MS technique presented as means of 6 biological replicates ± standard deviation. Statistically significant changes (p<0.05) are marked with asterisks.

#### Primary metabolite level in the transgenic HTZ lines

GC-MS technique allowed to determine the levels of soluble primary metabolites with sugars, organic acids and amino acids among them. In all transgenic lines a decrease of glucose, fructose and sucrose level was noted. The highest changes were observed for the line HTZ12 with decrease of 94%, 89,5% and 59,5%, respectively. In general, decreased levels of soluble sugar derivatives (sugar acids, sugar alcohols) were observed in all of the transgenic lines, except erythritol (elevated level in virtually all HTZ lines) (Supplementary File S2).

Among the primary metabolites organic acids play an important role as they participate in vital metabolic pathways, like glycolysis and Krebs cycle. The levels of these metabolites were found to be changed in the THZ transgenic lines. The most profound decreases were noted in all the transgenic lines for α-ketoglutarate (90% dropdown on average). The levels of citric acid and malic acid were also dropped down in all the transgenic lines but HTZ8 (Supplementary File S3).

Changes in free amino acids were also observed (Supplementary File S4). The highest decrease in the HTZ lines was found for proline and isoleucine. Leucine, valine, serine, asparagine and tryptophan levels dropped down to a lesser extent. HTZ12 line showed the most diversity from the control with arginine level increase of 3-fold and ornithine level of 8-fold.

#### Phenolic level in the transgenic HTZ lines

As catecholamine synthetic route is connected with the large pathway of phenylalanine transformation, the phenylpropanoid pathway, levels of particular phenolic derivatives (both free and cell wall bound) were determined by means of LC-MS technique. Based on retention times and absorption and mass spectra the presence of feruloyl-tyramine, kaempferol glycoside and chlorogenic acid was determined in the methanolic extracts (free phenolic fraction). In addition, based on spectral analysis, putative ferulic acid derivative and caffeic acid derivative were measured. The amount of feruloyl-tyramine was higher in all HTZ lines except HTZ43 compared to the control (1.66 ± μg/g DW) (Fig. 3). Similarly, the level of kaempferol glycoside was higher in all HTZ lines but HTZ43 in comparison to control (2.24 ± μg/g DW). Higher level of chlorogenic acid was found in HTZ8 and HTZ12 relatively to control (8.78 ± μg/g DW). The levels of the ferulic acid derivative and caffeic acid derivative was higher in all the transgenic lines (Supplementary File S5).

**Fig. 3.**
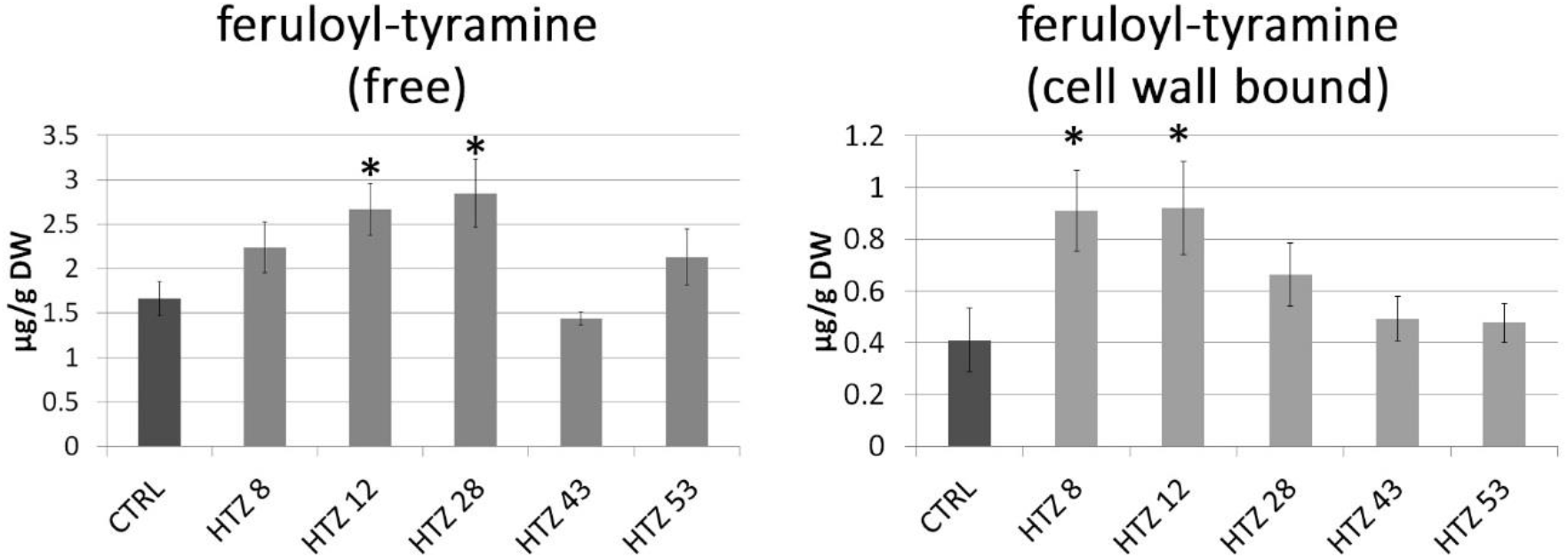
Amounts of free phenolic compounds in HTZ transgenic lines relatively to the control (CTRL – non-transgenic potato) obtained with LC-MS technique presented as means of 6 biological replicates ± standard deviation. Statistically significant changes (p<0.05) are marked with asterisks.

In the cell wall bound phenolic fraction 4-hydroxybenzoic acid (4HBA), 4-hydroxybenzaldehyde (4HBAl), vanillin, *p*-coumaric acid, ferulic acid and feruloyl-tyramine were measured. In addition, based on spectral analysis putative catechin derivative, *p*-coumaric acid derivative and tri-ferulic acid were identified. Levels of 4HBA and 4HBAl were elevated in all THZ lines (maximally 2-fold) in comparison to the control (0.71 μg/g DW and 0.32 μg/g DW, respectively). Similarly, vanillin level was increased in the HTZ lines (except HTZ43 – no change) relatively to control (0.76 μg/g DW). Levels of *p*-coumaric acid, ferulic acid and feruloyl-tyramine were most increased in HTZ8 and HTZ12, while insignificant changes were measured in the remaining transgenic lines (Fig. 3) (Supplementary File S5).

Total level of phenolic derivatives was higher in all the transgenic lines, both in methanolic extracts (free phenolics) and after hydrolysis (bound phenolics). The highest increase was noted for line HTZ28 and the smallest for line HTZ53 (Table 1).

**Table. 1.** Total phenolic content assayed in the transgenic HTZ lines in the methanolic extracts (free phenolics) and after hydrolysis (cell wall bound phenolics) compared to the control (CTRL).

|  | free phenolics [%] | cell wall bound phenolics [%] |
| --- | --- | --- |
| CTRL | 100 | 100 |
| HTZ8 | 212,3 | 148,5 |
| HTZ12 | 194,6 | 145,3 |
| HTZ28 | 249,7 | 169,4 |
| HTZ43 | 198,5 | 137,4 |
| HTZ53 | 132,5 | 107,5 |

### Antioxidant potential of the transgenic potato extracts

Phenolics are known to be strong antioxidants, thus their elevated level in the transgenic lines implies a higher antioxidant potential. The antioxidant potential of free phenolic extract and bound phenolic extract was measured using the DPPH method (Kedare and Singh, 2011). Methanol extracts from the transgenic plants were characterized with higher antioxidant potential (up to 5-times) compared to control (Fig. 4A). For cell wall hydrolysates improved antioxidant potential was noted for lines HTZ8 and HTZ12, but the changes were weaker (Fig. 4B).

**Fig. 4.**
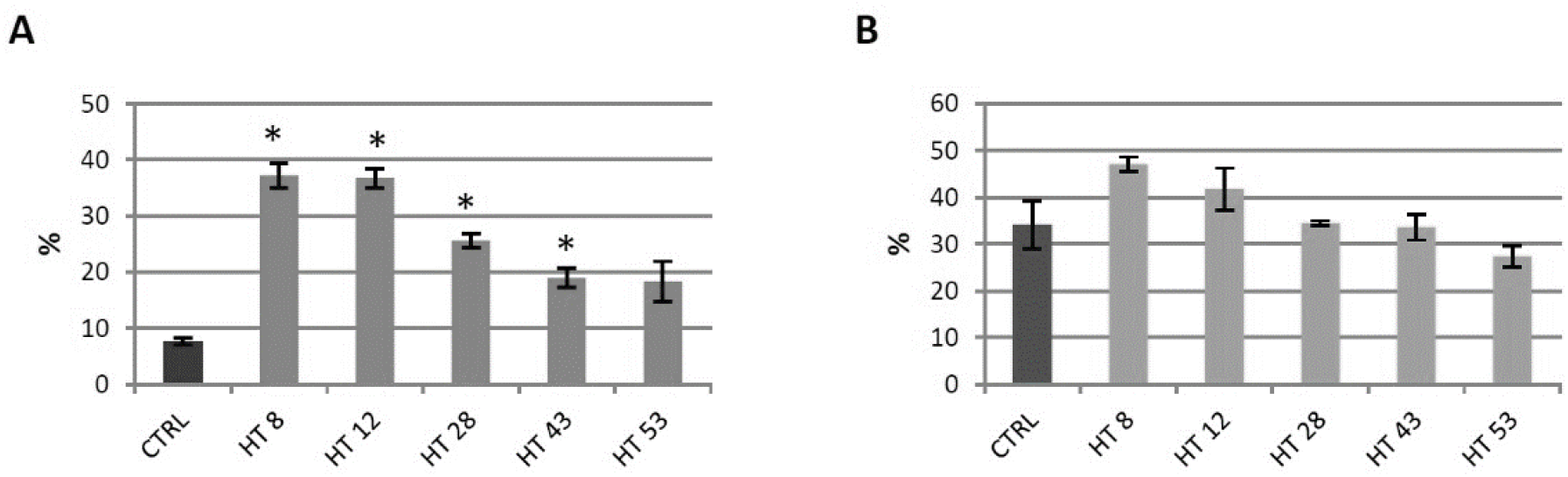
Antioxidant potential expressed as percent of quenched DPPH radical measured in methanol extracts (A) and extracts after hydrolysis (B) from transgenic HTZ lines compared to the non-transgenic potato (CTRL), presented as means of 6 biological replicates ± standard deviation. Statistically significant changes (p<0.05) are marked with asterisks.

### Red-ox analysis in the transgenic HTZ lines

#### Analysis of H_2_O_2_ level in transgenic potato plants

Level of H_2_O_2_, a gauge of the red-ox state was measured in the transgenic lines in relation to the control (Fig. 5). The level of H_2_O_2_ was elevated in all HTZ lines by 44% on average. The highest increase was measured in HTZ43 and HTZ53 (by 77% and 72%, respectively).

**Fig. 5.**
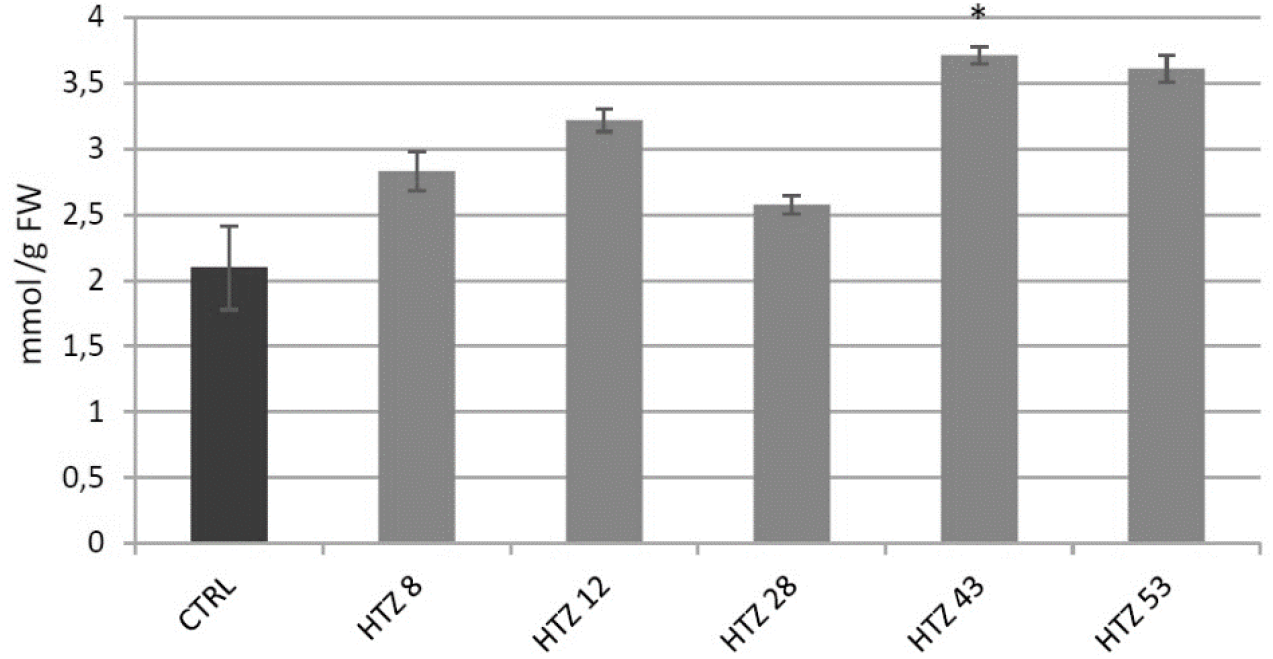
The level of H_2_O_2_ in HTZ potato lines compared to the non-transgenic potato (CTRL) presented as means of 6 biological replicates ± standard deviation. Statistically significant changes (p<0.05) are marked with asterisks.

#### Analysis of expression of genes involved in free radical processing in transgenic potato plants

The red-ox state depends mainly on the enzymatic system of free radical quenching, therefore the levels of transcripts of catalase (CAT), ascorbate peroxidase (APX) and three superoxide dismutase (SOD) genes were measured (Fig. 6). For catalase gene expression, a dropdown in all transgenic lines was noted (by 42% on average) with the highest changes for HTZ8 and HTZ12 (by 62% and 77%, respectively). Ascorbate peroxidase expression did not change in the transgenic lines, except HTZ53, where it reached 177% of the control. Three superoxide dismutase (iron, copper/zinc and manganese) gene expression levels were measured. For one of them – Cu/Zn SOD the average transcript level was considerably higher compared to the control (3.8-fold). The highest changes were detected for HTZ8 and HTZ12 (467% and 627% of the control, respectively). For Fe SOD and Mn SOD no significant changes were observed, except HTZ8, where a decrease (by 36% and 41%, respectively) was detected. In addition, a polyphenol oxidase (PPO) gene expression level was measured. The PPO is responsible for *o*-hydroxylation of monophenols to *o*-diphenols (catechols) and further to *o*-quinones. The expression level of PPO gene was substantially elevated in all the transgenic lines compared to the control (over 4 to 6 times).

**Fig. 6.**
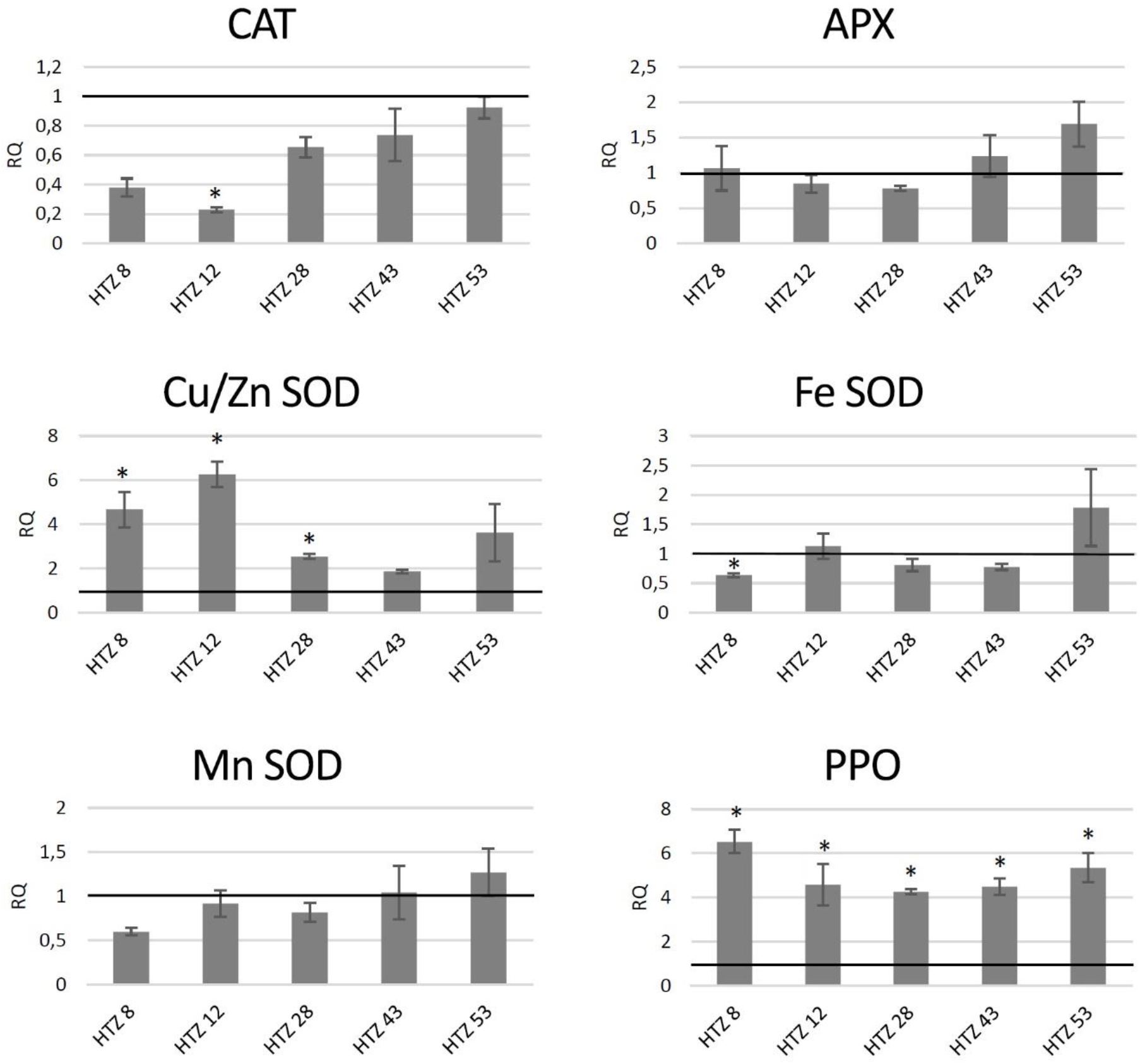
The level of transcript of genes involved in free radical processing in the HTZ lines presented as relative quantification (RQ) in relation to the non-transgenic control (horizontal line at RQ = 1). Elongation factor gene was used as a reference gene. The results were obtained with RT-qPCR method on cDNA matrix as mean values of 3 biological repeats ± standard deviation. Statistically significant changes (p<0.05) are marked with asterisks.

#### Analysis of H_2_O_2_ level in potato plants after exogenous L-DOPA treatment

Potato plants (wild type) from in vitro culture were treated with 0.1 mg/ml L-DOPA and then H_2_O_2_ level in 3, 6 and 12 hours later was measured (Fig. 7). Hydrogen peroxide level has increased in all three timepoints in comparison to the control, with the highest change at 3 hours after the treatment.

**Fig. 7.**
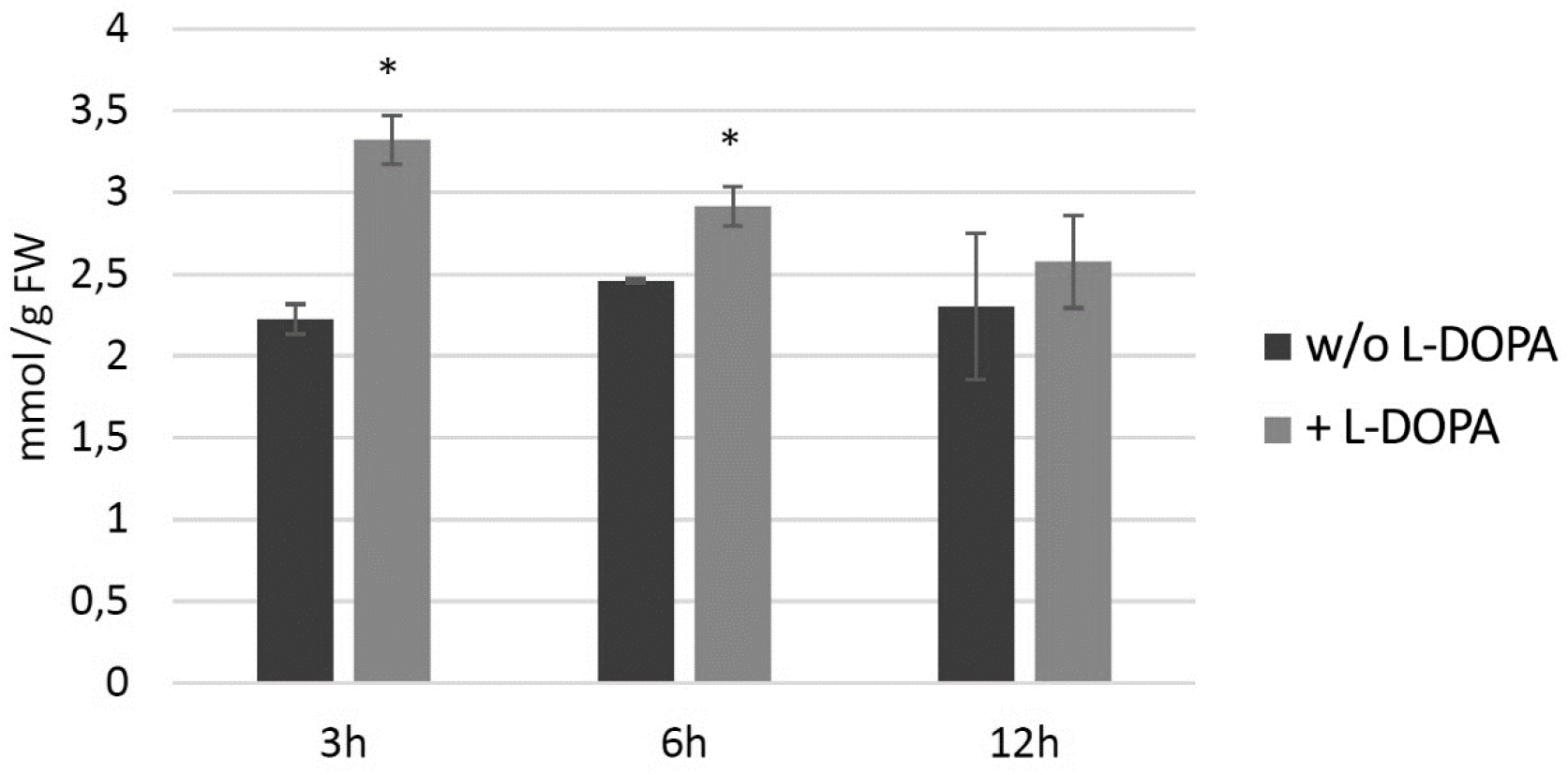
The level of H_2_O_2_ in wild type potato plants treated with 0.1 mg/ml L-DOPA (brighter bars) compared to the non-treated control (darker bars) presented as means of 6 biological replicates ± standard deviation. Statistically significant changes (p<0.05) are marked with asterisks.

#### Analysis of expression of genes involved in free radical processing in potato plants after exogenous L-DOPA treatment

In potato plants from in vitro culture treated with 0.1 mg/ml L-DOPA the levels of transcripts of genes involved in free radical processing were measured (Fig. 8). Transcript levels of genes connected with H_2_O_2_ neutralization was higher than in the control at 3 and 6 hours after L-DOPA treatment – catalase by ca. 30% and ascorbate peroxidase by ca. 45% and decreased below the control level at 12 hours after the treatment. Among the superoxide dismutase genes, expression of the one encoding the Cu/Zn SOD was higher at 3 and 6 hours after the treatment, by 74% and 94% compared to the control, respectively. Polyphenol oxidase gene was considerably activated (ca. 3-fold) at 6 and 12 hours after the treatment.

**Fig. 8.**
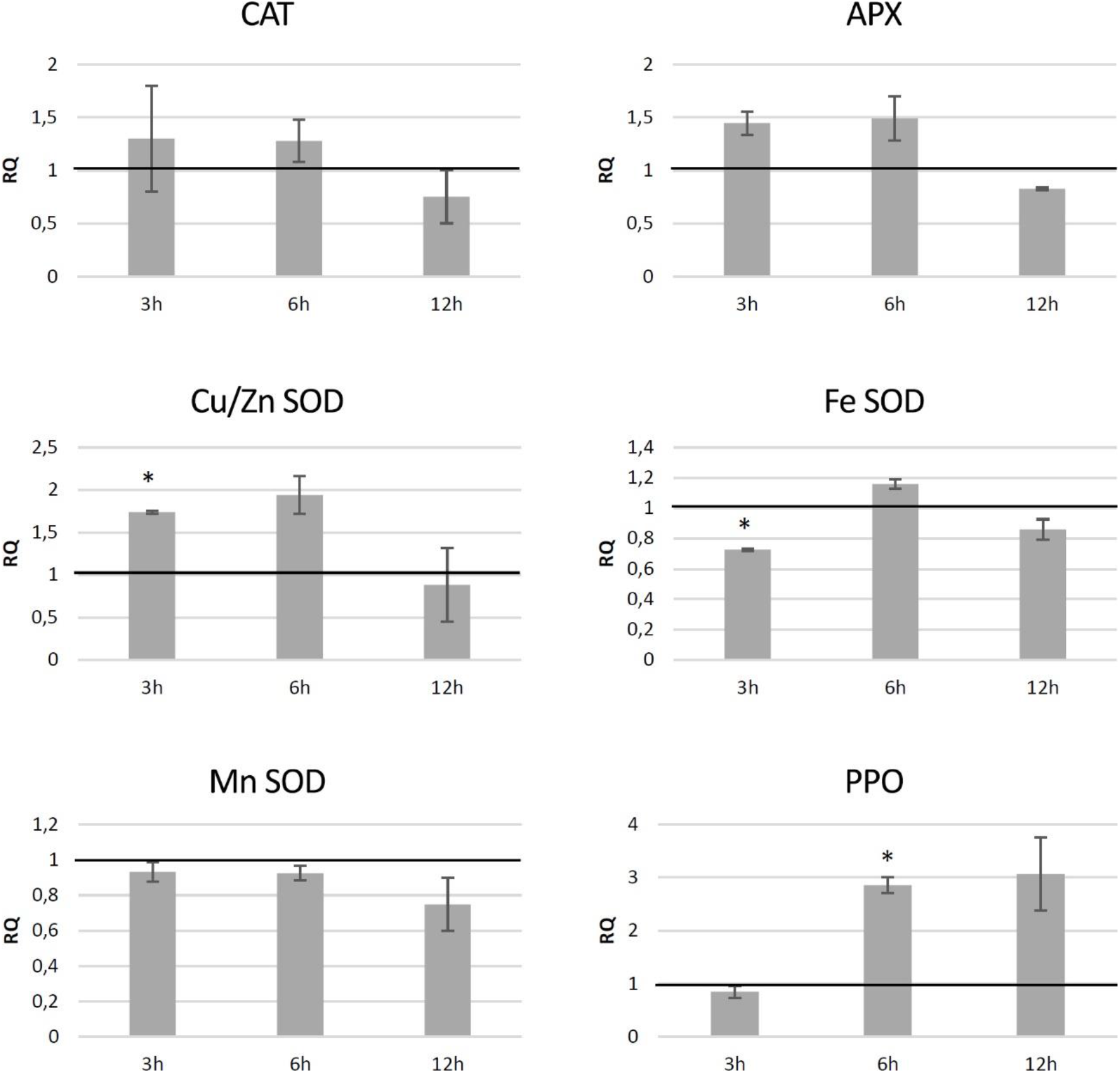
The level of transcript of genes involved in free radical processing in wild type potato plants treated with 0.1 mg/ml L-DOPA presented as relative quantification (RQ) in relation to the non-treated control (horizontal line at RQ = 1). Elongation factor gene was used as a reference gene. The results were obtained with RT-qPCR method on cDNA matrix as mean values of 3 biological repeats ± standard deviation. Statistically significant changes (p<0.05) are marked with asterisks.

### Resistance of the transgenic potatoes to Phytophtora infestans attack

It is suggested that catecholamines participate in plants’ response to stress, including pathogen attack (Świędrych et al., 2004). Moreover, the transgenic HTZ potato lines are characterized by higher antioxidant potential, which is known to positively correlate with resistance to pathogens (Kasote et al., 2015). In order to investigate the effect of expression of tyrosine hydroxylase on potato susceptibility to infection, the HTZ lines were submitted to infection with *Phytophtora infestans*, a common and dangerous potato pathogen. Lines HTZ12, HTZ28 and HTZ53 were the most resistant in comparison to the control (Fig. 9). No correlation was noted between the degree of infection and the phenolic compound level nor antioxidant potential.

**Fig. 9.**
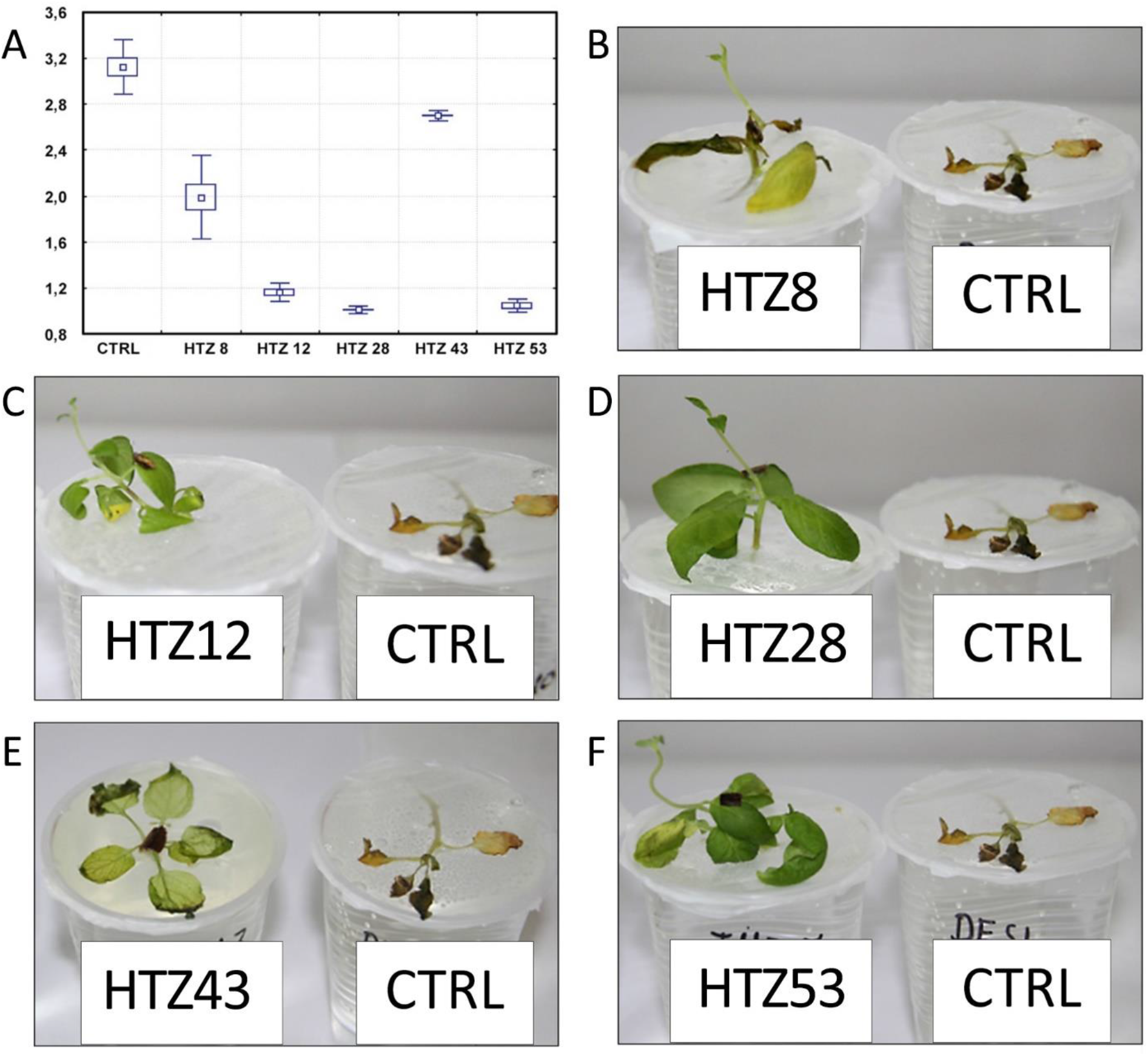
Degree of infection of transgenic HTZ lines with *P. infestans* in comparison to the non-transgenic potato (CTRL) (A). Sample photos of infected HTZ lines (B-E).

## Discussion

Although the literature on catecholamine biosynthesis in plants remains scanty, it is generally believed that the main pathway proceeds through decarboxylation of tyrosine to tyramine conducted by tyrosine decarboxylase (TD). The presence of the ‘alternative pathway’ involving hydroxylation of tyrosine to L-DOPA by tyrosine hydroxylase (TH), which is the main route of catecholamine production in animals, was not fully confirmed in plants, although L-DOPA, product of tyrosine hydroxylation, was found in these organisms. Based on the available plant genome sequence databases, no gene encoding tyrosine hydroxylase has been found. In this study, using transgenic potato plants with rat tyrosine hydroxylase gene, we analyzed the effect of the ‘alternative pathway’ of catecholamine synthesis on the plant’s metabolism. As anticipated, the obtained transgenic lines were characterized by increased L-DOPA levels, but despite further expectations we observed decreased levels of dopamine and subsequent compounds of this route. In light of the results obtained for transgenic potatoes with overexpression of tyrosine decarboxylase, where the catecholamine level was significantly increased (Swiedrych et al., 2004), the observed results suggest that the catecholamine biosynthesis route through tyrosine hydroxylation is non-functional in the potato plants, while the induction of the L-DOPA synthesis by heterologous tyrosine hydroxylase reorganizes catecholamine biosynthesis and leads to redirection of the substrate towards the production of other types of compounds.

It is known that L-DOPA is produced by some plants and utilized as allelochemical (Fujii, 2003, Nishihara et al., 2004) and its phytotoxicity is ascribed to the oxidative damage resulting from free radical species including ROS, generated from the pathway of L-DOPA metabolism (either autooxidation or oxidation aided by polyphenol oxidases (PPO)) to quinone and subsequently to melanins. Because in our research, after transformation of potato with tyrosine hydroxylase gene, we did not observe increased catecholamine synthesis, we suspected that the surplus of L-DOPA might have resulted in its oxidation followed by generation of free radicals and consequently in oxidative stress in the plants. Indeed, we have observed an elevated level of H2O2 in the HTZ transgenic potato lines, which may translate to a constant oxidative stress present in the plants. This stays in agreement with Arabidopsis response to L-DOPA treatment, which led to changes in transcriptional activities of genes associated with plant’s response to stress, both biotic and abiotic, with those involved in amino acid metabolism, oxidative stress, melanin synthesis and lignification among them (Golisz et al., 2010). Prolonged enhanced production of ROS can pose a threat to cells by causing the peroxidation of lipids, oxidation of proteins, damage to nucleic acids etc. (Hasanuzzaman et al., 2012), thus it is imperative for an organism to maintain equilibrium between ROS and antioxidative capacity. In order to cope with oxidative stress, plants have developed several enzymatic and non-enzymatic approaches to eliminate free ROS in cells. The increased content of H_2_O_2_ in the HTZ potato lines correlated positively with the activation of gene encoding the cytoplasmic superoxide dismutase (SOD). At the same time, catalase (*CAT*) gene transcript level was lowered. It may reflect the mechanism, in which transcription activity of *CAT* gene decreases in plants during stress (e.g. pathogen infection), perhaps due to the very H2O2 molecule itself (Yi et al., 2003) or accumulation of salicylic acid (Shim et al., 2003). This helps maintaining the H2O2 content at higher levels, which in turn translates into activation of many antimicrobial activities (Lamb and Dixon, 1997, Averyanov, 2009). The transgenic HTZ lines were also characterized with elevated levels of phenolic compounds and higher antioxidant potential. The phenolics comprise of several groups of compounds, like flavonoids, phenolic acids, lignin or benzoates and have a whole gamut of functions, including structural, antioxidant, signaling and regulatory. Expression of gene of phenylalanine ammonia lyase (*PAL*), which is considered a key enzyme of the phenolic synthesis, was increased in the transgenic HTZ lines. Bearing in mind a constant stress triggered by L-DOPA biosynthesis this is not surprising as *PAL* expression is known to increase upon ROS content increase and various stress conditions (Anna et al., 2011, Wada et al., 2014, Darmanti et al., 2018). Our observation confirms Soares et al. (2012) (Soares et al., 2012) report, where soybean which absorbed L-DOPA showed increased phenolic compounds, lignin content and the activities of related enzymes such as PAL and cinnamyl alcohol dehydrogenase (CAD). Although lignin content in the four-week-old plants from in vitro culture is barely determinable, we observed considerably high *CAD* gene and shikimate/quinate hydroxy-cinnamoyltransferase (*HCT*) gene transcript levels (Supplementary File S6). Increased expression of genes connected with lignification was also observed in L-DOPA treated *Arabidopsis* (Golisz et al., 2010). Stress conditions, especially pathogen infections coax plants to reinforce the cell wall with bio-reactive compounds to hinder microbial penetration into cells. Among the phenolics identified in the HTZ lines feruloyl-tyramine deserves special attention as on one hand it is a catecholamine derivative and on the other, it participates in plant’s response to pathogen infections (Jang et al., 2004), wounding (Pearce et al., 1998) and drought (Sprenger et al., 2016). The observed increase in the hydroxycinnamic acid-tyramine amide content in the transgenic lines may be somehow incomprehensible, because the level of tyramine, a direct substrate did not differ, nor was the tyramine decarboxylase gene (*TD*) expression higher compared to the wild-type control. However, it is conceivable that tyrosine acts as the substrate for both tyrosine decarboxylase, normally occurring in the plant and for tyrosine hydroxylase. Because of the oxidative stress elicited by L-DOPA, the produced tyramine is immediately used in the reaction of the hydroxycinnamic acid amide biosynthesis driven by tyramine n-hydroxycinnamoyl transferase (THT), while dopamine synthesis is reduced. *THT* gene encoding this enzyme is known to be activated in the hypersensitivity reaction, due to salicylic acid (SA) action (Conrath et al., 2006, Kalenahalli et al., 2015). Analysis of the promoter sequence of the potato *THT* gene revealed the presence of a number of elements interacting with WRKY and MYB transcription factors, known to be connected with oxidative stress response (Supriya et al., 2013, Ma and Constabel, 2019). Such elements were also found in the promoter sequences of THT from other plant organisms, like tomato (Etalo et al., 2013) or wheat (Kage et al., 2017). Moreover, such promoter elements are found in a number of genes involved in phenylpropanoid pathway, allowing for their control by SA in response to infection (Sarkar et al., 2018). WRKY among other transcription factor genes were shown to be activated in *Arabidopsis* upon L-DOPA treatment (Golisz et al., 2010). These resemblance to pathogen attack reaction in the HTZ potato lines might stand behind their better resistance to *P. infestans*.

Another explanation might be rerouting of tyramine to the hydroxycinnamic acid amide production due to the presence of tyrosine hydroxylase (TH). TH is a soluble enzyme occurring in cytoplasm, but it has been shown to interact with plasma membrane elements, like phosphatidylserine (Kuhar et al., 1999). In animal neurons TH is localized in the vicinity of the synaptic vesicle and mitochondria together with aromatic amino acid decarboxylase (AAD) and dopamine β-hydroxylase (DH), which, thanks to the availability of substrates can swiftly convert L-DOPA to dopamine and dopamine to norepinephrine, respectively (Wang et al., 2009). The co-localization of the three enzymes using immunoprecipitation approach in rats was also shown by Cartier et al. (Cartier et al., 2010). As for now, the presence of such or similar complexes that could involve tyrosine decarboxylase remains hypothetical. However, the presence of tyrosine hydroxylase in transgenic potatoes could lead to the rearrangement of the putative enzymatic complex and tyramine might be rerouted to the biosynthesis of hydroxycinnamic acid amides catalyzed by THT. Nonetheless, this hypothesis requires detailed research on the interactions between the enzymes of catecholamine biosynthesis in plants.

To summarize, the study on the effect of the alternative route of tyrosine metabolism towards catecholamines led to acquisition of the new knowledge on the possibility of manipulating plant metabolic pathways by introducing animal-type system. Expression of rat tyrosine hydroxylase gene in potatoes resulted in elevated L-DOPA level. However, this intermediate did not serve for catecholamine biosynthesis, but was oxidized with polyphenol oxidases and possibly also spontaneously. These reactions produced ROS molecules, which elevated level led to the formation of constant oxidative stress conditions. To compensate, antioxidant machinery was actuated, both enzymatic and non-enzymatic. This in turn caused better resistance of the transgenic potatoes to *P. infestans* infection. We suspect that introduction of an external element to the mechanism of catecholamine synthesis in potato disturbed it and caused redirection of their substrates. These results constitute a starting point to a new study, which will answer the question whether an enzymatic complex producing catecholamines exists or not.

## Materials and Methods

### Plant material

Potato plants *(Solanum tuberosum* L., cv. Desiree), obtained from “Saatzucht Fritz Lange KG” (Bad Schwartau, FRG), were grown in tissue culture on MS medium (Murashige and Skoog, 1962) supplemented with 0.8% agar, 1% sucrose and 250 mg/L claforan under 16 h light (21°C) and 8 h darkness (16°C) cycle.

### Construction of transgenic potato lines

The cDNA encoding TH from *Rattus norvegicus* (EMBL/GenBank database acc no. M10244.1), was ligated in the sense orientation into the plant binary vector under the control of the 35S CaMV promoter and OCS terminator. The vector was introduced into the *Agrobacterium tumefaciens* strain C58C1:pGV2260 and the integrity of the plasmid was verified by restriction enzyme analysis. Young leaves of wild-type potato *S. tuberosum* L. cv. Desiree were transformed with *A. tumefaciens* by immersing leaf explants in bacterial suspension. *A. tumefaciens* inoculated leaf explants were subsequently transferred to callus induction medium and shoot regeneration medium. Transgenic plants were pre-selected by using PCR with the primers for the neomycin phosphotransferase (kanamycin resistance) gene ((forward, CCGACCTGTCCGGTGCCC; reverse, CGCCACACCCAGCCGGCC)) on genomic DNA isolated from 3-week-old tissue-cultured plants as a template and then selected by northern blot analysis with a HT specific cDNA fragment as probe and subsequently by western blot analysis with antibodies specific to rat HT protein (Sigma-Aldrich).

### Northern blot analysis

Total RNA was prepared from frozen plant material using plant RNA purification kit (QIAGEN RNeasy Kit). Following electrophoresis (1.5%(_w/v_) agarose, 15%(_w/v_) formaldehyde), RNA was transferred to nylon membranes (Hybond N^+^, Amersham, UK). Membranes were hybridized overnight at 42°C in 250 mM sodium phosphate buffer (pH 7.2) containing 7%(_w/v_) SDS, 1%(_w/v_) bovine serum albumin (BSA) and 1 mM EDTA. Radioactively labelled full-length cDNA was used as a hybridization probe. Filters were washed three times in 1 × SSC containing 0.5%(w/v) SDS at 65°C (highly stringent condition) or in the same buffer but at 42°C (medium stringent condition) for 30 min.

### Western blot analysis

Proteins were extracted from frozen plant material using extraction buffer (100 mM Hepes-NaOH, pH 7.4, 10 mM MgCl2, 1 mM EDTA, 1 mM EGTA, 20%(v/v) glycerol, 0.5 mM PMSF, 70 mM beta-mercaptoethanol) supplemented either with 0.1% or 1%(_v/v_) TritonX-100. The assessment of the expression of rat TH gene by means of western blot analysis using rabbit IgG anti HT protein was conducted as described previously (Aksamit-Stachurska et al., 2008). Briefly, solubilized protein was run on 12%(_w/v_) SDS polyacrylamide gels and blotted electrophoretically onto nitrocellulose membranes (Schleicher and Schuell). Following transfer, the membrane was sequentially incubated with blocking buffer (5%(w/v) defatted dry milk), and then with antibody directed against the TH protein (1:5000 dilution). Formation and detection of immune complexes was performed with alkaline phosphatase-conjugated goat anti-rabbit IgG at a dilution of 1:1500.

### Isolation and analysis of phenolics

The frozen tissue (100 mg), comminuted in liquid nitrogen was extracted to 1 ml of 0.1% HCl in methanol in an ultrasonic bath for 10 minutes and centrifuged at 12000 × g, 4°C for 10 minutes. The procedure was repeated twice and the collected supernatants were merged, dried under nitrogen flow and re-suspended in 200 μl of methanol (free phenolic fraction). The remaining pellet was hydrolyzed in 2 M NaOH at 37°C overnight. Then the pH was adjusted to 3 using concentrated HCl and two volumes of ethyl acetate were added and mixed thoroughly. Following centrifugation (12000 × g, 4°C, 15 minutes), the organic phase was collected and the extraction to ethyl acetate was repeated two more times. All the collected volumes were merged, dried under nitrogen flow and resuspended in 200 μl of methanol (bound phenolic fraction). The obtained extracts were analyzed in UPLC with diode array detector 2996 PDA and mass detector QTOF. Acquity UPLC BEH C18 (2.1 × 100 mm, 1.7 μm) column was used for compound separation. The mobile phase consisted of 0.1%_(v/v)_ formic acid (solvent A) and acetonitrile (solvent B) and the flow was 95A:5B for 1 minute, then linearly for 11 minutes to 70A:30B, for next 4 minutes to 0A:100B and 1 minute to 95A:5B at rate of 0.4 ml/min. Column temperature was 25°C. Absorbance was measured at 210 nm – 500 nm range and mass spectra were registered in the range of 50-1000 Da under the following conditions: nitrogen flow 800 l/h, source temperature 70°C, cone temperature 400°C, capillary voltage 3.5 kV, cone voltage 40 V – 60 V, scan time 0.2 s (Wojtasik et al., 2013). Identification of compounds based on the comparison to the retention times, absorbance spectra and mass spectra of pure standards (Sigma-Aldrich).

### GC-MS analysis of metabolome

The amount of 150 mg of whole plant tissue ground in liquid nitrogen was extracted with methanol (12 ml/g FW). Samples were incubated at 70°C for 15 min. and centrifuged at room temperature (12000×g, 10 min.). Then they were supplemented with 1.5 ml H_2_O and extracted with 0.75 ml chloroform. The polar phase was dried and resolved in piridin/metoxyamine solution (20 mg/ml) and then incubated at 37°C for 2 hours. Samples were derivatized with N-methyl-N-trimethylsilyl trifluoroacetamide (MSTFA) and analyzed in GC-MS. GC8000 gas chromatograph with A2000 autosampler and Voyager Quadrupole Mass Spectrometer (Thermoquest) were used. Chromatograms and mass spectra were analyzed with TagFinder software based on mass spectra database (The Max Planck Institute of Molecular Plant Physiology (MPI-MP), Golm, Germany) (Roessner et al., 2000, Luedemann et al., 2008, Kopka et al., 2004). Ribitol was used as the external standard.

### RT-PCR

Total RNA prepared from frozen plant material using plant RNA purification kit (QIAGEN RNeasy Kit) was used for cDNA synthesis preceded by removal of residual genomic DNA with DNase I kit (Invitrogen) according to the producer’s instructions. High capacity cDNA Reverse Transcription Kit (Thermo Fisher Scientific) was used for reverse transcription of the RNA as stated in the producent’s protocol. Transcript levels were determined with RT-qPCR using StepOnePlus^™^ Real-Time OCR Systems (Applied Biosystems) on the cDNA matrix. Primers used in the reactions were designed in LightCycler^®^ Probe Design v2 (Roche) software. For the reactions DyNAmo SYBR Green qPCR Kit (Thermo Fisher Scientific) was used. The cDNA was dissolved 5 times prior to the experiment. The annealing temperature was 57°C. Changes in the transcript levels of the examined genes were presented as relative quantification in regard to control. Elongation factor gene was used as the reference gene.

### Determination of antioxidant capacity

Methanol extracts (free phenolic fraction and bound phenolic fraction) were used for antioxidant potential determination with 2,2-diphenyl-1-picrylhydrazyl (DPPH) (Kedare and Singh, 2011). 6 μl of sample extract was added to 200 μl of 0.2 mM DPPH methanolic solution and incubated at room temperature in darkness for 15 min. 6 μl of methanol was used in the control and pure methanol was used as blank. Subsequently, absorbance at λ=515 nm was measured. The antioxidant potential corresponded to the degree of free radical reduction and was presented as:

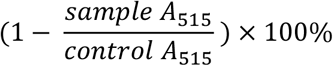

### Hydrogen peroxide assay

50 mg of four-week-old plant green tissue ground in liquid nitrogen were used for the analysis. Each sample was supplemented with 200 μl of 20 mM phosphate buffer pH 6.5, mixed and centrifuged (4°C, 12000×g, 10 min). 50 μl of the supernatant was then used for the assay with Amplex Red Assay (Life Technologies) according to the producer’s instruction. The absorbance was measured at λ=560 nm, fluorescence at λ_ex_ = 571 nm and λ_em_ = 585 nm (Wang et al., 2017).

### Treatment of plants with L-DOPA

Four-week-old potato plants from in vitro culture were submitted to 0.1 mg/ml L-DOPA treatment. The plants grown in glass jars (5 plants per a jar) were sprayed with 3 ml of L-DOPA and collected in 3, 6 and 12 hours after the treatment.

### Determination of resistance to pathogen

*Phytophtora infestans* (pathotype 33-IHAR PIB) was grown on solid PDA medium (3%) at 18°C for 14 days in darkness. Next, rye caryopses sterilized by autoclave, were placed onto the well grown mycelium (30 per a Petri dish) and were left for 7 days under the same conditions. The inoculum prepared in this manner was applied to a leave of individual plant. The plants were grown in hydroponic culture in a grow chamber (12 hour photoperiod with 15°C at day and 10°C at night) on liquid B5 medium (diluted 10 times) (Gamborg et al., 1968). The experiment was conducted for 14 days and the degree of infection was assessed according to a bonitation scale (1-1.5 – high resistance; 2-2.5 – mediocre resistance; 3-3.5 – weak resistance).

### Statistical analysis

All experiments were performed in at least 3 biological repeats. The results were presented as mean values ± standard deviation. For significance of changes evaluation one-way ANOVA with Tukey post-hoc or Kruskal-Wallis test was used (Statistica, Statsoft). Independent Component Analysis (ICA) for the data obtained from GC-MS analysis was performed with MetaGeneAnalyse software (v1.7.1; http://metagenealyse.mpimpgolm.mpg.de) (Scholz et al., 2005).

## Acknowledgements

This study was supported by grant No. 2017/24/C/NZ1/00393 from National Science Centre (NCN, Poland). We wish to thank to the group of Prof. Joachim Kopka (MPI Molecular Plant Physiology, Golm, Germany) for useful advices and great help in GC–MS experiments.

---

Supplementary File S1. Consecutive selection steps of potato plants transformed with HT gene from R. norvegicus. A – sample PCR product electrophoresis separation with use of primers specific to a 486-nt fragment of HT cDNA from R. norvegicus. P – positive control, N – negative control (non-transgenic potato); B – sample Northern blot of RNA samples from transgenic potato plants bearing HT gene from *R. norvegicus*. P – positive control, N – negative control (wt potato); C – Western blot analysis of proteins isolated from transgenic HT potato plants. N – negative control (non-transgenic potato).

Supplementary File S2. Levels of soluble sugars in HTZ transgenic lines relatively to the control (non-transgenic potato) obtained with GC-MS technique presented as means of 6 biological replicates ± standard deviation. Statistically significant changes (p<0.05) are marked with asterisks.

Supplementary File S3. Levels of organic acids in HTZ transgenic lines relatively to the control (non-transgenic potato) obtained with GC-MS technique presented as means of 6 biological replicates ± standard deviation. Statistically significant changes (p<0.05) are marked with asterisks.

Supplementary File S4. Levels of soluble amino acids in HTZ transgenic lines relatively to the control (non-transgenic potato) obtained with GC-MS technique presented as means of 6 biological replicates ± standard deviation. Statistically significant changes (p<0.05) are marked with red color.

Supplementary File S5. Levels of soluble (A) and cell wall bound (B) phenolics’ derivatives in HTZ transgenic lines relatively to the control (non-transgenic potato) obtained with LC-MS technique presented as means of 6 biological replicates ± standard deviation. Statistically significant changes (p<0.05) are marked with red color.

Supplementary File S6. The levels of transcripts of cinnamyl-alcohol dehydrogenase (CAD) and shikimate/quinate hydroxy-cinnamoyl transferase (HCT) genes in the HTZ lines presented as relative quantification (RQ) in relation to the control (horizontal line at RQ = 1). Elongation factor gene was used as a reference gene. The results were obtained with RT-qPCR method on cDNA matrix as mean values of 3 biological repeats ± standard deviation. Statistically significant changes (p<0.05) are marked with asterisks.

